# Facilitated Dissociation of Transcription Factors from Single DNA Binding Sites

**DOI:** 10.1101/135947

**Authors:** Ramsey I. Kamar, Edward J. Banigan, Aykut Erbas, Rebecca D. Giuntoli, Monica Olvera de la Cruz, Reid C. Johnson, John F. Marko

## Abstract

The binding of transcription factors (TFs) to DNA controls most aspects of cellular function, making the understanding of their binding kinetics imperative. The standard description of bimolecular interactions posits TF off-rates are independent of TF concentration in solution. However, recent observations have revealed that proteins in solution can accelerate the dissociation of DNA-bound proteins. To study the molecular basis of facilitated dissociation (FD), we have used single-molecule imaging to measure dissociation kinetics of Fis, a key *E. coli* TF and major bacterial nucleoid protein, from single dsDNA binding sites. We observe a strong FD effect characterized by an exchange rate ∽1 × 10^4^ M^-1^s^-1^, establishing that FD of Fis occurs at the single-binding-site level, and we find that the off-rate saturates at large Fis concentrations in solution. While spontaneous (i.e., competitor-free) dissociation shows a strong salt dependence, we find that facilitated dissociation depends only weakly on salt. These results are quantitatively explained by a model in which partially dissociated bound proteins are susceptible to invasion by competitor proteins in solution. We also report FD of NHP6A, a yeast TF whose structure differs significantly from Fis. We further perform molecular dynamics simulations, which indicate that FD can occur for molecules that interact far more weakly than those we have studied. Taken together, our results indicate that FD is a general mechanism assisting in the local removal of TFs from their binding sites and does not necessarily require cooperativity, clustering, or binding site overlap.

**SIGNIFICANCE STATEMENT:** Transcription factors (TFs) control biological processes by binding and unbinding to DNA. Therefore it is crucial to understand the mechanisms that affect TF binding kinetics. Recent studies challenge the standard picture of TF binding kinetics by demonstrating cases of proteins in solution accelerating TF dissociation rates through a facilitated dissociation (FD) process. Our study shows that FD can occur at the level of single binding sites, without the action of large protein clusters or long DNA segments. Our results quantitatively support a model of FD in which competitor proteins invade partially dissociated states of DNA-bound TFs. FD is expected to be a general mechanism for modulating gene expression by altering the occupancy of TFs on the genome.

**Author Contributions:** Ramsey I. Kamar

designed research, performed research, contributed new reagents/analytic tools, analyzed data, wrote the paper

Edward J. Banigan

Aykut Erbas

Rebecca D. Giuntoli

Monica Olvera de la Cruz

designed research, performed research, wrote the paper

Reid C. Johnson

John F. Marko

## INTRODUCTION

Protein-DNA interactions ultimately control all aspects of cellular function through their actions as “transcription factors” (TFs) by regulating gene transcription, folding DNA into chromosomes, and modifying the structure of chromatin; these regulatory and structural functions are often interwoven (1-9). Understanding protein-DNA interaction kinetics is therefore essential to the mechanistic understanding of cellular function. The standard picture of protein-DNA interactions assumes binding via concentration-dependent association and concentration-independent “spontaneous dissociation” kinetics, with net affinity neatly described by the ratio of the off-rate, *k*_off_, to theassociation rate constant, *γ* , *i.e.*, *K*_D_ = *k*_off_ / *γ*. Experiments that resolve dynamics of individual molecules are starting to challenge this classical picture: lifetimes of DNA-protein complexes have been found to be appreciably shortened by nearby proteins that compete for space on DNA (10-20). However, the molecular mechanisms underlying this “facilitated dissociation” (FD) effect remain unclear.

A well-characterized DNA-binding protein that has shown strong FD effects is the *E. coli* TF Fis (13, 14). This dimeric TF binds diverse but sequence-specific DNA sites through a pair of helix-turn-helix domains to form a stable protein-DNA complexes (21). Fis also has a weaker, but appreciable, non-sequence-specific DNA-binding affinity, that contributes to its role as a chromosome-organizing protein (22). This previous body of work establishes the Fis-DNA complex as a model system for studying the molecular basis of FD.

In this paper we present single-molecule experiments on the Fis-DNA system plus theoretical analyses which establish that, through FD, competitor proteins can strongly modulate the stability of a single protein dimer bound to a single DNA binding site, leading to a concentration-dependent off rate. The off-rate saturates at high protein concentration, indicating the presence of a rate-limiting step along the FD pathway. Our experiments establish that FD can occur at the single-binding-site level without the need for cooperative effects via clusters of multiple proteins or long segments of DNA (13). We also find that spontaneous dissociation and FD have distinct dependences on salt concentration. Our data for Fis are globally and quantitatively described by an analytically tractable theory in which FD is generated by partial dissociation of the initially bound Fis, thereby allowing an unstable Fis-DNA-Fis ternary complex to form (10, 11, 13, 16, 18-20, 23-28). We validate our analytical theory using molecular dynamics simulations of FD that explicitly incorporate ionic effects. Finally, additional experimental data for a monomeric yeast TF, NHP6A, which binds a different short DNA through a single HMG box interaction (9), also displays FD, indicating that the two DNA-binding domains of Fis are not required for a protein-DNA complex to display concentration-dependent dissociation kinetics. Based on our results, we expect FD to be a generic effect that modulates the effective affinity of TF-DNA interactions in cells.

## RESULTS

### Fis stably binds a short minimal binding sequence

We used single-molecule fluorescence imaging to measure the dissociation kinetics of gfpFis from individual DNA binding sites (Fig. 1). We immobilized one end of 27-bp Cy3-labeled F1 dsDNAs to cover slips; the high affinity F1 Fis binding sequence has been well characterized thermodynamically and structurally (21). While Fis is able to bind to a core 21 bp region of this site (21), additional contacts over the 27 bp window stabilize binding (29, 30), and shorter DNA oligos demonstrate weaker binding effects (21, 25, 30), indicating 27 bp to be the complete binding site length.

**Figure 1.**
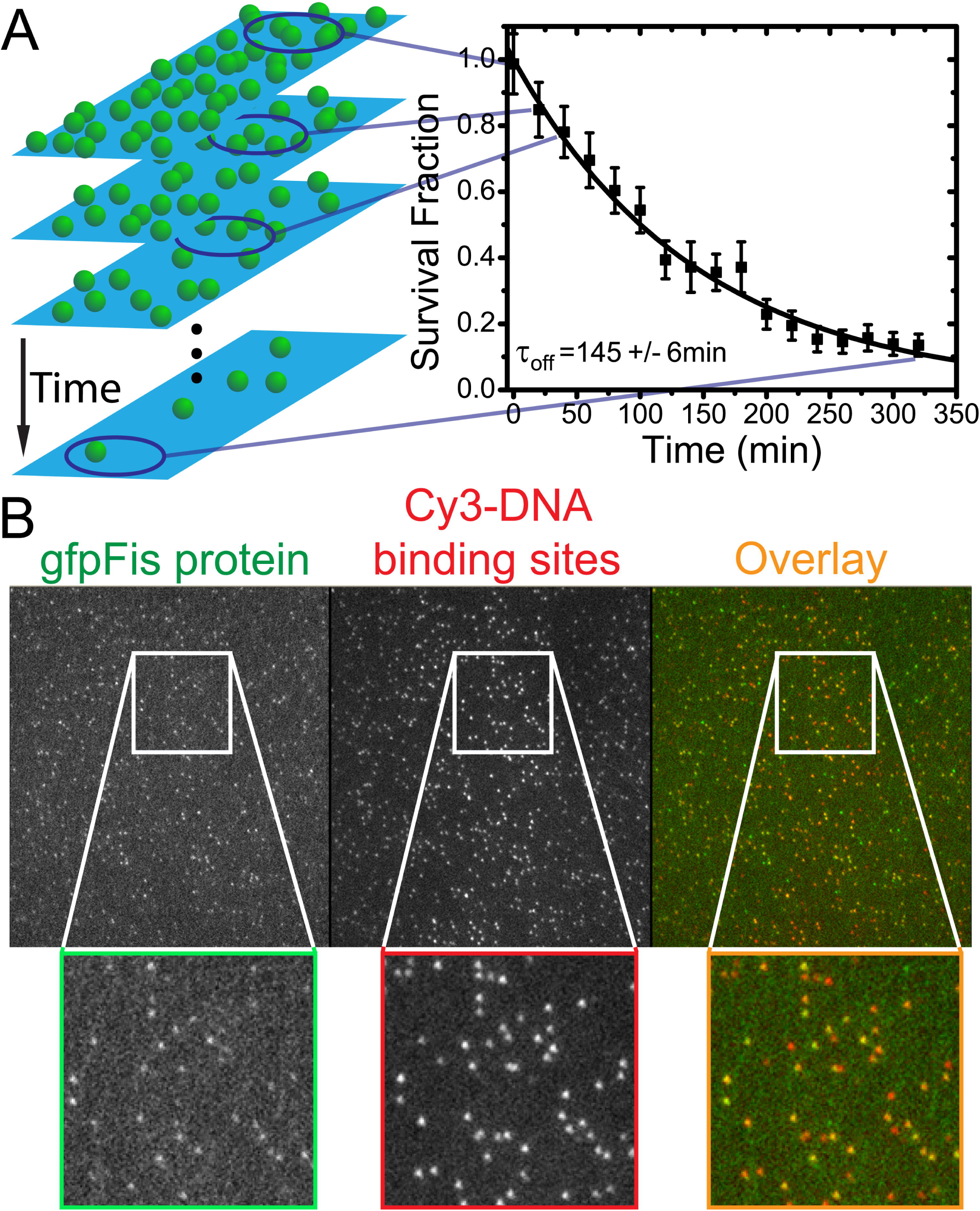
Off-rate measurement. (*A*) The survival fraction is measured by counting the number of fluorescent signals (green spheres, left) remaining in the flow cell as a function of time and normalizing by the initial number of signals. A new region along the flow cell is used for a measurement of the survival fraction at each subsequent time point. To obtain the off-rate, the survival fraction decay is fit to a single decaying exponential exp(-*t*/*τ*_off_) (right). In example shown, *τ*_off_ = 145 ± 6 min with *χ*^2^/*ν* = 0.34. See *SI Appendix* for details of survival fraction calculation. (*B*)Camera frame showing single molecule fluorescence image in separate channels. Each panel is the full 512X512 pixel array (52.5X52.5 µm^2^). Insets are magnified views of the region contained in the white box. Fluorescent signals from gfpFis (left panel) and signals from Cy3-labeled F1 DNA binding sites (middle panel) show a high degree of colocalization (right panel) indicating the specificity of gfpFis binding to F1 sequences. In the right panel, gfpFis signals are false colored green and Cy3-DNA signals are false colored red. Regions where green false color overlaps with red false color appear orange indicating colocalization. Only gfpFis signals that co-localize with Cy3-DNA signals are retained for inclusion in the measurement of survival fraction.

We determined the off-rate of gfpFis in protein-free, 100 mM NaCl buffer by measuring the number of gfpFis molecules that remain bound to F1 DNAs as a function of time (Fig. 1*A*). We observed a high degree of colocalization (up to ∽85%) between signals in the gfpFis and DNA channels (Fig. 1*B*), which ensures that the GFP signals we retain for analysis correspond to Fis dimers bound to DNA. The decay curves fit well to single exponential decays of the form exp(–*t*/**τ**_off_) ; asample decay curve is shown in (Fig. 1*A*). In competitor-free buffer, Fis remains stably bound for along period (**τ**_off_ = 180 ± 40 min, 3 replicate experiments) giving a spontaneous off-rate of 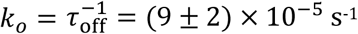.

### Fis protein in solution accelerates the off-rate of Fis from single binding sites

Because previous experiments demonstrating FD for Fis involved protein initially bound along a long extended dsDNA (13), potentially containing overlapping Fis binding sites or multi-protein clusters, we were interested in whether FD could also occur at a single F1 binding site. We first confirmed that F1 sequences only allow for the stable binding of individual gfpFis dimers by recording gfpFis signal fluorescence trajectories in protein-free buffer and constructing histograms of the number of bleaching steps (Figs. S1*A* and *B*). We observe that the majority (≈94 %) of trajectories bleach in one or two steps as expected for single gfpFis dimers (*SI Appendix*, Fig. S1).

We next recorded a series of gfpFis decay curves measured with different concentrations of wtFis in solution (0-1790 nM, Fig. 2*A*). Prior to adding wtFis, the excess gfpFis was washed out of the flow cell. We observed that with increasing Fis concentration the off-rate curves decay increasingly rapidly, demonstrating that FD of Fis does not require long segments of DNA that contain multiple binding sites. The decay curves fit well to single exponential decays, and the resulting off-rate (*k*_off_ =1/**τ**_off_, the rate constant from the exponential fit), shows an initially linear increase with protein concentration (Fig. 2*B*). However, for wtFis concentrations beyond ∽250 nM the off-rate saturates. This indicates the presence of a rate-limiting step in the protein dissociation pathway (25).

**Figure 2.**
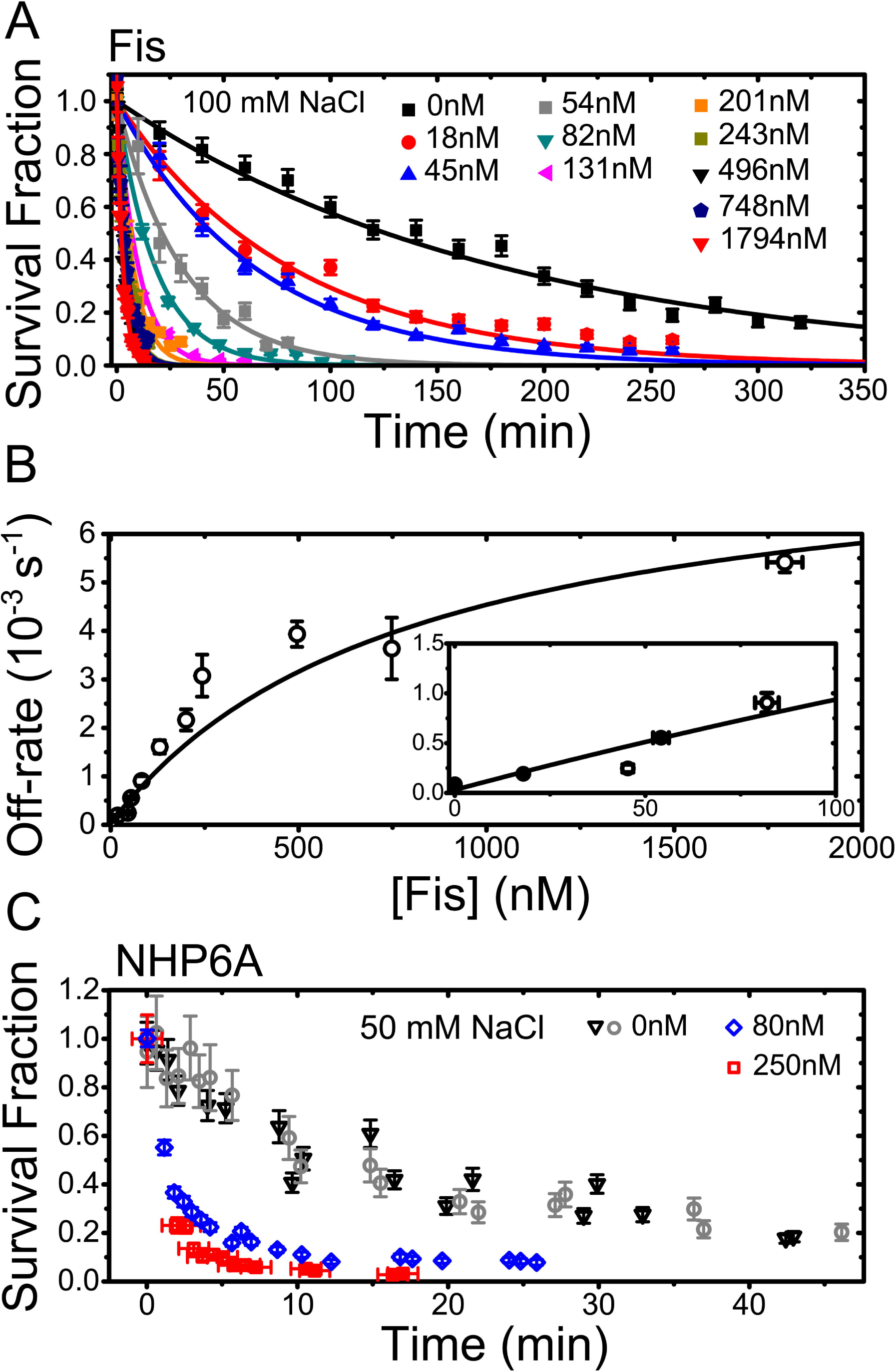
TF dissociation measurements from single binding sites. (*A*) Sample survival fraction time course measurements (solid symbols) for gfpFis at each concentration of wtFis in solution that was tested. Error bars are estimates of the statistical uncertainty in the data points from various sources (*SI Appendix*). At each concentration, the early portion of the survival fraction decay is well fit to a single exponential decay to obtain the off rate (solid curves, typically *χ*^2^/*ν* ∽1). (*B*) Off-rate versus concentration of wtFis in solution. Vertical error bars are a weighted SD of 2-4 measurements, except for the measurement at 54 nM which contains a single measurement (*SI Appendix*). Sizes of the horizontal error bars are smaller than the symbols (except for data point at 1794 nM) and represent statistical error in [wtFis]. Inset: Low-concentration behavior. Solid curve is a fit to Eq. 1. The exchange rate *k*_exch_ and saturation rate *k_sat_* are estimated from the fit and are given by *B*^-1^-*DA*/*B*^2^ = (1.0 ± 0.2) × 10^4^ M^-1^ s^-1^ and *A*^-1^ = (8.1 ± 1.5) × 10^-3^s^-1^, respectively; *D* is within 0-8 nM,*χ*^2^/ ≈ 9. Errors are scaled by.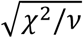 If the ratio *D*/*B* is fixed to the measured value (*D*/*B* =*k*_o_ = (9 ± 2) × 10^-5^ s^-1^), *B* and *A* change by 16.9% and 7.6%, respectively, which are within error. (*C*) NHP6A survival fraction time courses showing that NHP6Agfp also displays FD from single binding sites using wtNHP6A as a competitor. Experiments performed in protein-free, 50mM NaCl buffer are shown for two duplicate trials (blue and black symbols). Vertical error bars are estimated as in (*A*). The data sets corresponding to 80 nM and 250 nM [wtNHP6A] are normalized to the number of signals measured after the survival fraction has already decayed by re-incubating the flow cell with NHP6Agfp and counting the number of signals under protein-free buffer conditions.

### Fis off-kinetics are described by a simple model with a ternary intermediate

Our results can be described using the kinetic scheme depicted in Eq. S6 and Fig. S2*A* (13, 27, 31) which specifies a possible mechanism by which TFs in solution lead to FD of a TF bound to DNA. This model is based on the idea that there are thermally excited, *partially dissociated* states where some, but not all, DNA-protein contacts are broken. A TF fully bound to its binding site (state 0) is thermally excited into a partially bound state (state 1) that is susceptible to invasion by a TF from solution to form an unstable ternary complex (state 2). The off-rate of Fis corresponds to the inverse of the mean time 〈**τ**_off_〉 for afully-bound Fis molecule to transition to a fully-unbound state (state 3), either by spontaneously dissociating (transitioning directly from state 1 to state 3) or by FD.

A calculation (Eq. S10 and *SI Appendix*) of the mean time to dissociation under this scheme gives the following form for the off-rate (25):

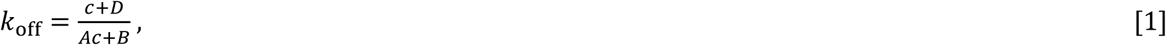

where *c* is [wtFis]. The constants *A*, *B*, and *D* are combinations of the microscopic rate constants from the kinetic model. By using Eq. 1 to fit the data (Fig. 2*B*), we determine 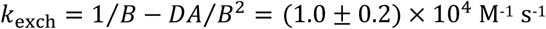 The saturated rate at high wtFis concentration is 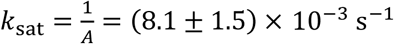 which is nearly 100-fold larger than the spontaneous dissociation rate *k*_o_ = (9 ± 2) × 10^-5^ s^-1^. Qualitatively, our observation that the off-rate of Fis from DNA is accelerated by Fis proteins in solution, and quantitatively, the approximate exchange rate, are both in agreement with previous single-DNA, protein competition experiments (13) as well as with experiments demonstrating FD for Fis bound to the *E. coli* nucleoid (14). Our results demonstrate that FD does not necessarily require long stretches of DNA that contain overlapping binding sites (32) or clusters of proteins.

### NHP6A also displays FD from SRY binding sites

To determine whether dimeric structure (e.g. Fis) is required for FD, we measured survival probability decay curves of NHP6A, a monomeric TF in yeast that consists of a single HMGB box domain (9), using the same approach we used for Fis. For a NHP6A binding site, we used a short Cy3-labelled dsDNA segment containing the recognition sequence for the SRY protein since an NMR solution structure for the SRY DNA-NHP6A complex has been determined (9). Survival probability curves of NHP6Agfp fusions, measured in 50mM NaCl buffer that includes 80nM or 250nM wtNHP6A, showed faster decays than those in protein-free buffer (Fig. 2*C*). Our results with monomeric NHP6A, which has a different structure and binding mode than dimeric Fis, demonstrate the generality of FDS

### Salt dependence of spontaneous off-rate is strong; salt dependence of FD is weak

Returning to the case of Fis, we measured the univalent-salt-concentration dependence of the off-rate. We reasoned that this would provide a molecular level probe of the kinetic pathways involved in FD. We first measured the salt dependence of the off-rate in protein-free buffer (Fig. 3*A* and *C*), at several salt concentrations, *cc*S, in the range 75-250 mM NaCl, and we observed a strong salt dependence: the off-rate fits to a power law of the form 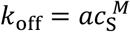, with exponent *M* = 2.6 ± 0.3. The overall salt dependence of the spontaneous dissociation pathway (0 → 1 → 3 in Fig. S2*A*) is therefore strongly salt-dependent.

**Figure 3.**
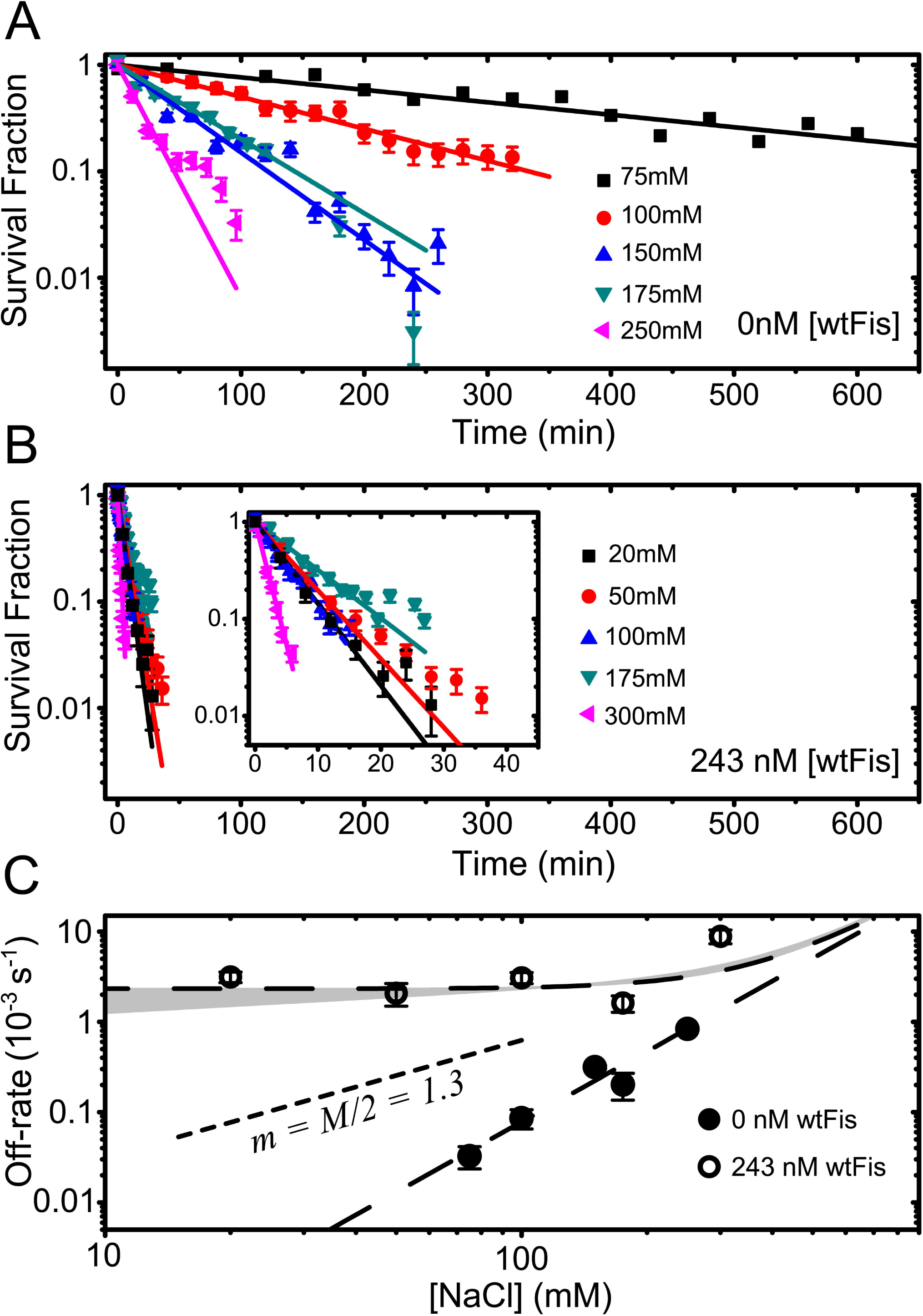
Salt dependence of off-rate. (*A*) gfpFis decay curves measured in protein-free buffer at multiple NaCl concentrations. (*B*) gfpFis decay curves measured in buffer containing 243 nM [wtFis] at multiple NaCl concentrations. Inset: Same data shown on a zoomed in scale to show detail. In both (*A*) and (*B*), the early portion of the survival fraction curves are fit to a single exponential decay to obtain the dissociation rate, and the error bars are estimated just like in Fig. 2*A*. (*C*) Off-rate of gfpFis as a function of NaCl concentration in protein-free buffer (solid symbols) and buffer containing 243 nM [wtFis] (open symbols). Error bars are estimated as in Fig. 2*B*. Long-dashed curves are powerSlaw fits. The protein-free off-rate is fit to a single power-law 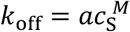 giving *M* = 2.6 ± 0.3. The 243 nM off-rate is fit to a sum of power-laws (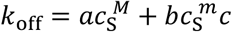, Eq. 2) with *M* and *a* fixed to thevalues obtained by fitting the protein-free data. Grey band is a range of fits to the 243 nM data obtained by allowing joint variation in *m* and *b* while still allowing for a plausible fit. In these fits *mm* ranged from 0 to 0.25, beyond which the slope of the power-law did not allow for reasonable agreement with the data. A single power-law exponent equal to *M*/2 is shown for comparison (short-dashed curve).

To probe the [wtFis]-dependent FD pathway, we next measured the off-rate with 243 ± 6 nM wtFis in solution at several *cc*S in the range 20-300 mM NaCl (Fig. 3*B* and *C*; this wtFis concentration is roughly halfway up the FD off-rate curve of Fig. 2B, and is also likely comparable to the free wtFis concentration found *in vivo* (14)). The observed salt dependence is weak (Fig. 3*C*), suggesting the protein-dependent pathway either does not depend on salt concentration or that it is rate-limited by a salt-independent step. This suggests fitting the 243 nM data to the combined power-law form

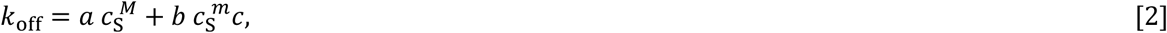

where *a* and *M* are fixed to the values obtained by fitting the protein-free data. The fits to the 243 nM data (Fig. 3*C*) can accommodate a power-law exponent no larger than *m* ≈ 0.25 with the best fit given by *m* = 0 (i.e., no salt dependence at all).

It is well-established that the dissociation constant for many protein-DNA interactions has a power-law dependence on univalent salt concentration, *c*_*S*_, leading to log *K*_*D*_ vs. log *c*_*S*_ being linear (33, 34):

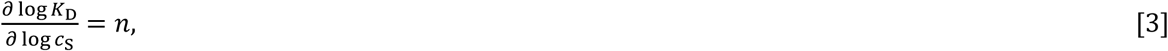

where the slope *n* is proportional to the number of positive counterions released from the DNA molecule, or, equivalently, the number of contacts formed between the protein and the DNA when the protein binds (35). This motivates the use of power laws in Eq. 2 to fit the salt dependence of the off-rate in Fig. 2C since *k*_off_ = γ*K*_D_ for a simple bimolecular reaction. It should be noted that the partial dissociation model makes the prediction (which we have confirmed here, Fig. 3*C*) that over some range of *c*_S_, which depends on the protein concentration, the salt dependence of the off-rate is weaker when proteins are in solution (depicted in Fig. S2*B* and explained in *SI Appendix*). These results suggest that the number of counterions released in forming a ternary complex during FD is a small fraction of the number of counterions that condense on the binding site when Fis is released during spontaneous dissociation.

### Partial unbinding model simultaneously explains *c*_*s*_- and *c*-dependence

We sought to generalize Eq. 1 to account for the observed salt dependence. We formulated a model (Fig. 4*A*) that incorporates the effects of DNA-bound counterions on the free energies involved in the kinetic steps along both dissociation pathways. The model also allows for asymmetric unbinding of the protein from the DNA. We utilized the exact analytical calculation (above) of the mean time 〈**τ**_off_〉 (Eqs. 1 and S10) for a protein to dissociate from the DNA binding site in terms of the microscopic rate constants *kij* (arrows in Fig. 4*A*) and the protein concentration, *c*=[wtFis], in solution. Using detailed balance, the salt dependencies of the set of *kij* are included, which results in a model for the off-rate, *k*_off_ =〈**τ**_off_〉^-1^, in terms of the physical parameters of the model, the protein concentration, and the ionic strength (see *SI Appendix* for detailed derivation). The incoming wtFis molecule forms a ternary complex by binding to the partially exposed binding site, destabilizing the binding of the original gfpFis molecule, and blocking its return to state 1. In our model, this leads to the dissociation of both the initially bound gfpFis and the incoming wtFis molecules (arrow *k*_23_).however it is also possible that the wtFis molecule replaces the original molecule (arrow *k*_20_*).

**Figure 4.**
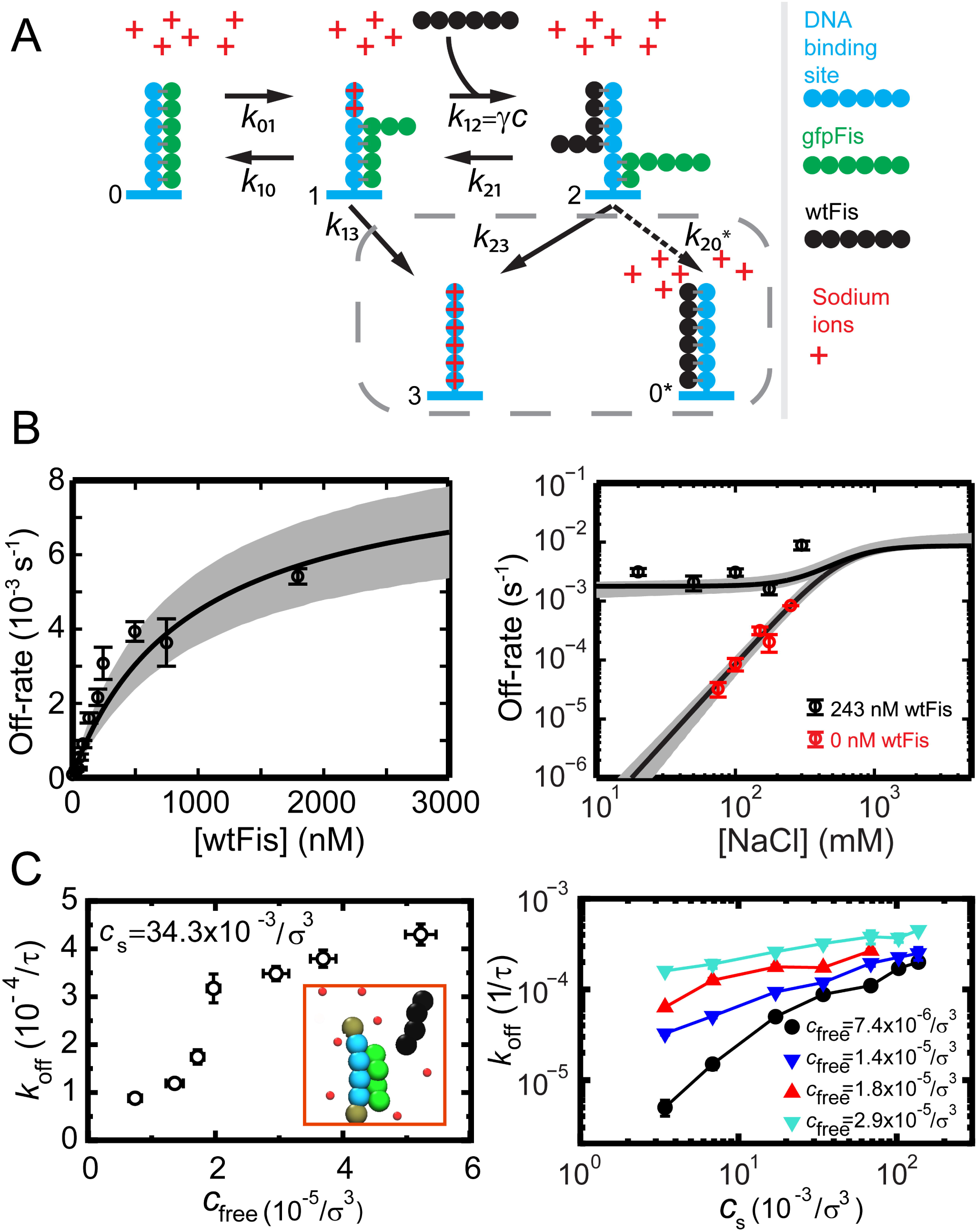
Global fit to kinetic model and simulations. (*A*) Schematic representation of a kinetic model of facilitated dissociation. Schematic depicts the multivalency of Fis-DNA interactions by drawing Fis as a multipartite object. However, this should not be interpreted as Fis taking a linear form. Model explicitly includes positive Na+ ions in solution, which can condense on DNA and compete with binding locations on Fis for contacts to the DNA. Contacts made between gfpFis and the DNA are represented by grey bars. Going from state 1 to 2, the original TF (green) is shown with fewer contacts to the DNA to depict the possibility that the competitor could destabilize the binding of the original TF. Grey box encircles the two possible final states of the ternary complex, however this study considers the left-pointing solid arrow. See *SI Appendix* for detailed derivation of the mean reaction time including the salt and protein concentration dependence with this kinetic model. (*B*) [wtFis] (left panel) and [NaCl] (right panel) dependence measurements of the off-rate are globally fit to extended kinetic model (solid curves). Bimolecular on-rate constant ;γ = 1.04 ± 0.19 × 10^8^ M^-^1^^s^-^1^^ is fixed by experiment (*SI Appendix*) in the fitting. Grey bands represent 68.3% confidence intervals of the fit (*SI Appendix*). (*C*) Coarse-grained simulations corresponding to extended kinetic model where protein and DNA molecules are represented by chains of reactive beads as illustrated in the inset (*SI Appendix*). Left panel: protein concentration dependence of the off-rate obtained from the simulations. Off-rate is plotted in units of the inverse self-diffusion time **τ** of a bead (*SI Appendix*). Right panel: salt concentration dependence of the off-rate at multiple protein concentrations.

We performed a global fit to the *c*- and *c*_s_-dependences of the off-rate using this model (see *SI Appendix* for fitting details). Since the large number of parameters made fitting impossible, we fixed the bimolecular on-rate constant *γ* by direct measurement (Fig. S4 and *SI Appendix*) to reduce the number of free parameters. We made the simplifying assumption that the measured on-rate constant *γ* meas corresponds to the rate *γ* for Fis to bind to sites that already contain partially bound Fis molecules in state 1. However, we expect that *γ* as defined in the model is somewhat less than*γ*meas since it is measured on initially empty binding sites. A smaller;*γ* will lead to fitted values for the microscopic kinetic constants that differ from those that come from our fit to the model (Table 1).However, the fit is well constrained by the data and is shown in Fig 4*B*. Furthermore, the kinetic rate *k*_01_ is well constrained by the data, even before *γ* is fixed. All the kinetic rates of the model, except *k*_23_, which has a broad distribution (Table 1), are well constrained by the data. However, it is clear that *k*_23_ is large compared to *k*_01_, so *k*_01_ is rate limiting for the competitor dependent pathway. Despite the fact that the microscopic rate constants (except for *k*_01_) that come from the fit cannot albe obtained accurately, the model is able to simultaneously capture the weak (essentially absent) salt dependence of the 243 nM data between 20-175 mM NaCl, the strong salt dependence of the protein-free measurements, and the saturation of the off-rate at high protein concentrations. Therefore our findings support partial unbinding of Fis as the molecular mechanism of FD.

**Table 1.** Microscopic rate constants obtained from global fitting with FD model.*

| Rate constant | Constraints from fitting or measurement† |
| --- | --- |
| $k_{01}$ | $8.7^{+1.4}_{-1.4} \times 10^{-3} \text{ s}^{-1}$ |
| $k_{10}$ | $90^{+30}_{-20} \text{ s}^{-1}$ |
| $k_{13}$ | $900^{+310}_{-270} \times 10^{-3} \text{ s}^{-1}$ |
| $k_{23}$ | $4^{+24}_{-2} \times 10^6 \text{ s}^{-1}$ |
| $k_{21}‡$ | $k_{21} \leq k_{23}/600 \quad k_{21} \sim k_{23}/10^{9 \pm 1}$ |
| $\gamma$ | $1.04^{+0.19}_{-0.19} \times 10^8 \text{ M}^{-1} \text{ s}^{-1}$ |
| $n_{01}$ | 0-0.25 |
| $n_{13}$ | $2.55^{+0.30}_{-0.25}$ |

\* Rate constants are referenced to their values at 100mM NaCl. $\gamma$ , $n_{01}$ , and $n_{13}$ are not fitted parameters, but are instead measured values whose distributions are imposed on the fit in Fig. 4B.
† Except for $k_{21}$ , all constraints are statistical 68% confidence intervals. Quoted values are those that minimize $\chi^2$ with $\gamma$ , $n_{01}$ , and $n_{13}$ fixed to their measured, most probable values. Additional systematic uncertainty in the kinetic rates comes from $\gamma$ being known only up to an overall unknown factor. Note the combination of $n_{01} + n_{13}$ , and not $n_{13}$ directly, is measured. Uncertainty on $n_{13}$ is therefore estimated from direct measurement of $n_{01}$ and by simulating errors on $n_{13}$ given the experimental errors on $n_{01}$ and $M = n_{01} + n_{13}$ .
‡ $k_{21}$ was not a fitting parameter in the final fitting function Eq. S21 ( $k_{21}$ is ignored in comparison to $k_{23}$ in Eq. S19), however placing constraints on it was possible by performing fits using an equation derived from Eq. S19. The upper limit on $k_{21}$ is found by fixing $\alpha_{21}$ to $\frac{\alpha_{23(100 \text{ mM})} n_{13} - n_{01}}{N}$ and finding the minimum value of $N$ that allows consistency with the data.

### Coarse-grained simulations validate multivalent partial unbinding model

To judge the applicability of the multivalent partial unbinding model, we performed coarse-grained MD simulations (Fig. 4*C* and *SI Appendix*) designed to mimic features of our experiment and our revised model by a generic bead spring-model (see the inset of Fig. 4*C*). Specifically, the simulations explicitly included multivalent structure in the DNA binding sites and competitor protein molecules, and, in addition, explicitly included positive charges in solution that could serve as counterions to compete for binding with positively charged protein molecules. The individual DNA binding sites are each represented by a chain of four negatively charged beads, and the proteins are represented by chains of four positively charged beads. All beads interact via long-and short-range Coulomb interactions. The steric interactions between beads are taken into account by repulsive excluded volume potentials; attractive interactions are only between the binding site and proteins (SI Appendix). We note that while we define our model via interaction potentials, it is possible to directly implement chemical reaction rates instead (36).

The simulations show a strongly accelerated off-rate of pre-bound protein when competitor proteins are introduced (Fig. 4*C*, left), qualitatively in accord with both our experimental data and our analytical model. The simulations also display the saturation of off-rate at large competitor concentrations (Fig. 4*C*, left) seen experimentally and in the analytical model. Finally, the simulations also produce a strong salt dependence in the absence of protein in solution (Fig. 4*C*, right, black curve) and weak salt dependence at high protein concentrations, again qualitatively in accord with experiment and the analytical model.

It is important to note that for reasons of computational power, the MD simulations must be done for weaker and consequently shorter-lived protein-DNA interactions than occur for Fis or NHP6A in our experiments. In the simulations, the length scale σ which corresponds to the size of a bead and is set by the requirement that the electrostatic energy between two beads in surface contact be equal to *k*_B_*T*, is approximately 0.7 nm, making 1/σ^3^≈5 M and 1/**τ**≈4x10^9^ s^-1^, where **τ** is the self-diffusion time of a bead, *k*_B_ is Boltzmann’s constant, and *T* is the absolute temperature, indicating that comparison of MD and experiment can be only qualitative. However, the MD results indicate that similar FD effects can be expected for biomolecule interactions that are far weaker and shorter-lived than we can study with our current time resolution.

## DISCUSSION

### Kinetic survival fraction measurements are well suited for studying the molecular basis of FD

In this study, we have shown that the TF Fis undergoes facilitated dissociation from single binding sites formed from short DNAs. Previous single DNA, protein competition experiments on Fis demonstrated FD on extended 48.5kb DNA molecules (13). However, interpretation of these experiments is complicated by the fact that there are multiple binding sites of varying affinity on long DNA molecules, which can lead to non-exponential decays of survival probability (27). Moreover, binding sites that overlap could assist FD (32) or allow for other types of cooperative dissociation events. This initially raised the question of whether FD with Fis requires protein clusters, the action of overlapping binding sites, or whether the competitor acts by sliding along adjacent DNA segments. Since our study uses a single binding site with a single binding strength, our results indicate that FD with Fis does not require long DNAs or cooperative binding effects, and this is further supported by our observations of single-exponential decays (Figs. 1*A*, 2*A*, and 3*A*-*B*).

Since we can readily measure slow off rates over a large dynamic range of TF concentration (Fig. 2), our method of measuring the survival fraction is able to observe FD of very high affinity TFs (*K*_D_ ≈ 1 pM for Fis-F1 interaction at low [wtFis] in our experiment). We are able to observe saturation of the off rate which, until now, has only been observed in DNA competition experiments (25). In contrast, detection of FD on high affinity TFs, displaying slow off-rates, might be missed using methods that rely on the collection of on/off binding events (e.g., smFRET), due to fluorophore bleaching hampering long observation times and limitations on the concentration of competitor that can be used (high competitor concentrations would lead to intervals between binding events that are shorter than the time resolution). This likely explains why FD of NHP6A was not seen using smFRET in a previous study (37).

### Microscopic picture of FD with Fis

By fitting our results to an exactly solvable theoretical model including salt effects, we developed a microscopic picture of FD (Fig. 4*A*). At high wtFis concentration, the off-rate saturates at *k*_sat_ = *k*_01_*k*_23_/(*k*_01_+*k*_23_) ≈ *k*_01_ (from fitting with the model, we find *k*_23_ *≫ k*_01_), indicating that the rate-limiting step along the FD pathway is the rate at which a fully bound protein is thermally excited into a partially bound state, thereby exposing the binding site to invasion by a competitor. There is a waiting time of ∽2 min (given by 1/*k*_sat_) for a bound Fis molecule to become susceptible to invasion, after which the formed ternary complex quickly leads to dissociation. That *k*_23_ is large compared to *k*_13_ demonstrates that the original TF is more strongly bound in the partially dissociated state (state 1) than in the ternary complex (state 2). The transition from state 1 to 3 is essential for the model to capture the spontaneous dissociation of Fis at low competitor concentration.

The nearly absent salt dependence of the FD pathway reflects the weak salt dependence of the rate-limiting kinetic step from states 0 → 1 (Figs. 3*C* and 4*A*), and suggests that few protein-DNA contacts are broken along the pathway leading to a ternary complex. It also indicates that during FD, the two Fis molecules are within one screening length relative to each other so that the total number of protein-DNA contacts is approximately conserved by the formation of the ternary complex. While a rigorous quantitative understanding of the microscopic picture of FD with Fis is limited by our knowledge of the bimolecular on-rate *γ*, our analytic model qualitatively describes our data remarkably well.

Additionally, we note that DNA near the boundaries of the 27 bp binding site may well be involved in FD, and thus, further single-molecule studies of slightly shorter or longer DNA oligos or of different flanking sequences might be useful in further understanding Fis-F1 FD. It is also unclear the degree to which protein conformational fluctuations play a role in FD. While relatively unstructured peptides have a plausible mechanism as in the case of e.g., polymerase-processivity clamp interactions (38), it remains to be understood how fluctuations of more stably folded proteins such as Fis or NHP6A allow FD to occur. Intriguingly, there is a hint in the data of a more complex (sigmoidal) shape to the rate-concentration curve (Fig. 2B) which may be indicative of multiple intermediate states along the FD pathway (24). Molecular dynamics simulations may be able to provide some hints of dynamical mechanism (20), although accessing the very long time scales associated with FD of Fis will be extremely challenging even with a coarse-grained model.

A previous study has described how competitors in solution can accelerate dissociation of molecular complexes by occluding rapid rebinding events and has demonstrated accelerated DNA duplex dissociation by this mechanism (39). However, the concentration scale, *c*, for this effect to occur is given by *c* ≈ 1/*δ*^3^, where *δ* is the size of the reaction volume, which in our case is of the order ∽10 nm. This would suggest a concentration scale of *c* ≈ 2 mM for FD, yet we already see a significant enhancement in off-rate at 20 nM. In addition, for a binder and target that are roughly the same size, inhibition of rebinding typically leads to only an approximately two-fold effect since the expected number of rebinding events is of order 1, but we observe a nearly 100-fold effect. Finally, it is hard to explain the observed weak salt dependence of the protein-dependent pathway using this type of model since the intermediate state involves a fully unbound protein. Taken together, our results argue against a rapid rebinding model and support the formation of a ternary complex as the mechanism underlying the concentration dependent off-rate of Fis. This conclusion is further supported by recent computational studies exploring the energy landscape of Fis binding that show the existence of a Fis-DNA-Fis ternary complex (20) and our simulations showing that FD via a ternary complex mechanism qualitatively recapitulates our experimental observations (Fig 4*C*).

### Generic nature of FD

We expect that FD should occur for any TF-DNA interaction that involves multiple contacts between the TF and DNA as long as the TF-DNA complex can partially unbind and expose the binding site to invasion by competitors. Our observation of FD of a monomeric protein, NHP6A, supports this assertion (Fig. 2C). We also note that we have observed heterotypic FD (Fis-driven dissociation of NHP6A, Fig. S7, see also (13)), which is indicative of a rather generic mechanism.

FD is unmistakable in experiments like ours, and those of others (11-13, 15, 16, 18), which have single-molecule dynamics that are observable at ≈1s time scales due to strong protein-DNA interactions (*K*_D_ ≤ 100 nM). However, FD is likely a general effect, controlling the unbinding kinetics of proteins with µM affinities, and typical single-molecule experiments with > 10 ms time resolutions are unable to observe these sub-ms dynamics. Our MD simulations, which by necessity use weak binding sites, support this assertion, exhibiting FD at high competitor concentrations. This suggests that FD could occur *in vivo* for typical TFs that bind much more weakly to DNA than Fis.Furthermore, even though a high concentration of an individual type of weakly binding TF may not be present *in vivo*, high concentrations of other proteins can be expected, and it has already been seen that competitor TFs of one type can cause FD of another type (13, 14). Therefore, it is reasonable to expect that FD could be an important mechanism for buffering the effective *K*D and assisting in the local exchange of a large class of TFs from chromatin in the nucleus.

### Physiological relevance of FD

In E-coli, it is observed that Fis is largely replaced by other nucleoid-associated proteins during slow bacterial growth (40). Our observation that competitors accelerate the dissociation of Fis from 1 pM affinity F1 sites suggests that FD plays an important role in this exchange and possibly serves as a mechanism to modulate the occupancy of strong Fis binding sites at high protein concentrations *in vivo*. Indeed, it has been observed that Fis binding lifetimes on the nucleoid are faster *in vivo* (where free Fis concentrations are at least several hundred nM) than they are on isolated nucleoids (14, 41), which suggests FD driven by a cytoplasmic concentration of Fis (and other DNA-binding proteins) in the few hundred nM range.

Our results suggest FD could have a profound effect on the dynamics of biological processes that depend on the binding of TFs *in vivo*. Cellular gene expression profiles and protein concentration levels occurring in complex regulatory networks should be affected by FD through its ability to shorten the residency time of a wide class of TFs that control these networks. FD may also play a role in regulating the dynamics of chromatin structure by facilitating the exchange of TFs with other regulatory proteins such as histones, nucleosomes, and remodelers from chromatin (6, 8, 42). In particular, FD could be a mechanism for regulating the ability of high affinity TFs to switch transcription on and off, and possibly facilitates TF mobility along the genome. Furthermore, our simulations suggest that FD should accelerate removal of proteins that bind DNA far more weakly than Fis (Fig. 4C) at timescales far shorter than our current single-molecule experiment can access. In conclusion, FD of TFs may be an important general effect to take into account in systems biology simulations that model gene expression in cells.

## MATERIALS AND METHODS

Single-molecule experiments were carried out using Cy3-and biotin-labeled 27 bp double-stranded DNA oligomers which were attached to the interior of a flow cell via streptavidin and biotin-PEG. Proteins were introduced via flow, including gfp-labeled proteins. The DNAs and gfp-labeled proteins were imaged using total-internal-reflection fluorescence (TIRF) imaging which allowed individual DNAs and proteins to be observed. Time series of protein-occupation of DNAs were collected and then analyzed to obtain protein binding kinetics. Coarse-grained molecular dynamics (MD) simulations were carried out using a simulation box containing 100 sparsely placed surface-grafted semiflexible DNA chains (binding sites), an equal number of protein (flexible) chains initially bound onto the DNA chains, a prescribed number of initially unbound proteins, counterions for protein and DNA chains and prescribed number of monovalent salt ions. The DNA and proteins are modeled by a coarse-grained bead-spring model with both short and long-range electrostatic interactions. Further details of single-molecule and simulation methods may be found in the *SI Text*.

## DATA SHARING STATEMENT

All data, documentation, and code used in analysis will be made available on request to the communicating author.

## ACKNOWLEDGMENTS

Work at NU was supported by the NIH through grants R01-GM105847, U54-CA193419 (CR-PS-OC) and a subcontract to grant U54-DK107980, and by the NSF through grants MCB-1022117, DMR-1611076 and DMR-1206868. Work at UCLA was supported by NIGMS grant GM038509.

